# Variant calling from scRNA-seq data allows the assessment of cellular identity in patient-derived cell lines

**DOI:** 10.1101/2021.04.13.439634

**Authors:** Daniele Ramazzotti, Fabrizio Angaroni, Davide Maspero, Gianluca Ascolani, Isabella Castiglioni, Rocco Piazza, Marco Antoniotti, Alex Graudenzi

## Abstract

**Matters Arising from:** Sharma, A., Cao, E.Y., Kumar, V. et al. Longitudinal single-cell RNA sequencing of patient-derived primary cells reveals drug-induced infidelity in stem cell hierarchy. *Nat Commun* **9**, 4931 (2018). https://doi.org/10.1038/s41467-018-07261-3.

In Sharma, A. et al. *Nat Commun* **9**, 4931 (2018) the authors employ longitudinal single-cell transcriptomic data from patient-derived primary and metastatic oral squamous cell carcinomas cell lines, to investigate possible divergent modes of chemo-resistance in tumor cell subpopulations. We integrated the analyses presented in the manuscript, by performing variant calling from scRNA-seq data via GATK Best Practices. As a main result, we show that an extremely high number of singlenucleotide variants representative of the identity of a specific patient is unexpectedly found in the scRNA-seq data of the cell line derived from a second patient, and vice versa. This finding likely suggests the existence of a sample swap, thus jeopardizing some of the translational conclusions of the article. Our results prove the efficacy of a joint analysis of the genotypic and transcriptomic identity of single-cells.

---

The integration of omics data from single-cell sequencing experiments enables the analysis of cell-to-cell heterogeneity at unprecedented resolution and on multiple levels [3]. This is especially relevant in the study of cancer evolution and will be essential to shed light on the key mechanisms underlying intra-tumor heterogeneity, metastasis, drug resistance and relapse [4]. In particular, scRNA-seq experiments are increasingly employed, typically in the characterization of the gene expression patterns of single-cells in a variety of experimental settings [6]. However, an increasing number of studies is proving that scRNA-seq data can be used to efficiently call genomic variants, thus providing an available and cost-effective alternative to whole-genome/exome and targeted sequencing [5, 16]. Despite known pitfalls, such as the impossibility of calling variants from non-transcribed regions and the typically high rates of noise and dropouts [9], the mutational profiles so obtained can be promptly used to determine the identity of single-cells. This aspect is important, for instance, in the analysis of the clonal evolution of tumors and in the detection of rare clones [11].

In [13], the authors employ single-cell transcriptomic data from patient-derived primary and metastatic oral squamous cell carcinomas (OSCC) cell lines (from a previously characterized panel [1]), to investigate possible divergent modes of chemo-resistance in tumor cell subpopulations. We integrated the analyses presented in the manuscript, by performing variant calling from scRNA-seq data via GATK Best Practices [2]. On the one hand, this analysis may allow one to reconstruct the longitudinal evolution of the tumor in presence of the treatment [10]. On the other hand, this allows one to deliver an explicit mapping between genotype and phenotype of single cells, thus providing important hints on the relation between clonal evolution and phenotypic plasticity. This might have a significant translational relevance, given the current shortage of accurate and affordable technologies for DNA and RNA sequencing of the same cells, despite the recent introduction of new protocols [7, 8].

In particular, we selected the scRNA-seq datasets of two cell lines derived from distinct OSCC patients –HN120 and HN137 – which include different data points, marked with the following suffixes: -P (primary line), -M (metastastic line), -CR (after cisplatin treatment), -CRDH (after drug-holiday). Since for the HN137P cell line two library layouts are provided (*single-end* and *paired-end*), which we here consider separately, and no HN137MCRDH is provided, we have a total of 12 datasets (all datasets are included in the GEO online repository, accession code GSE117872; please refer to the original article [13] for further details on the experimental setup).

In detail, we selected single cells labeled as “good data” on the GEO repository e performed variant calling (the whole procedure is described in detail in the Supplementary Material). 4, 924, 559 unique variants were detected on a total of 1, 116 single cells included in all datasets. Given the known limitations and the high levels of experimental noise of scRNA-seq data, we then applied a number of quality-control filters on variants, to ensure high confidence to the calls and to reduce the number of both false alleles and miscalls. In particular, we *removed*: (*i*) indels and other structural variants – to limit the impact of possible sequencing and alignment artifacts, (*ii*) variants mapped on mitochondrial genes, (*iii*) variants on positions with total read counts < 5 in > 50% of the cells in each time point – to focus the analysis on well-covered positions, (*iv*) variants detected in less than 20% of both HN120P and HN137P (*single-end*) cells – to focus on recurrent variants, (*v*) variants detected (≥ 3 alternative reads) in *both* HN120P and HN137P (*single-end*) – to define a list of variants that clearly characterize the identity of the two primary cell lines. We finally selected the variants observed in at least 1 cell (≥ 3 alternative allele reads, ≥ 5 total reads) of HN120P and in exactly 0 cells of HN137P (*single-end*), *and* the variants observed in at least 1 cell (≥ 3 alternative allele reads, ≥ 5 total reads) of a HN137P (*single-end*) and in exactly 0 cells of HN120P.

As a result, we identified 67 single-nucleotide variants (SNVs) that are representative of HN120P cell identity, and that are present in 0 cells of HN137P (*single-end*). Such variants are observed in high frequency in HN120P *and* in HN137P (*paired-end*), HN137PCR, HN137PCRDH, HN137M, HN137MCR, whereas are not observed (< 1% of the cells) in HN120PCR, HN120PCRDH, HN120M, HN120MCR, HN120MCRDH and HN137P (*single-end*). In Figure 1A we display the mutational profiles of all single-cells in all datasets. The total allele reads matrix and the alternative allele reads matrix for such variants are provided as Supplementary Table 1.

**Figure 1:**
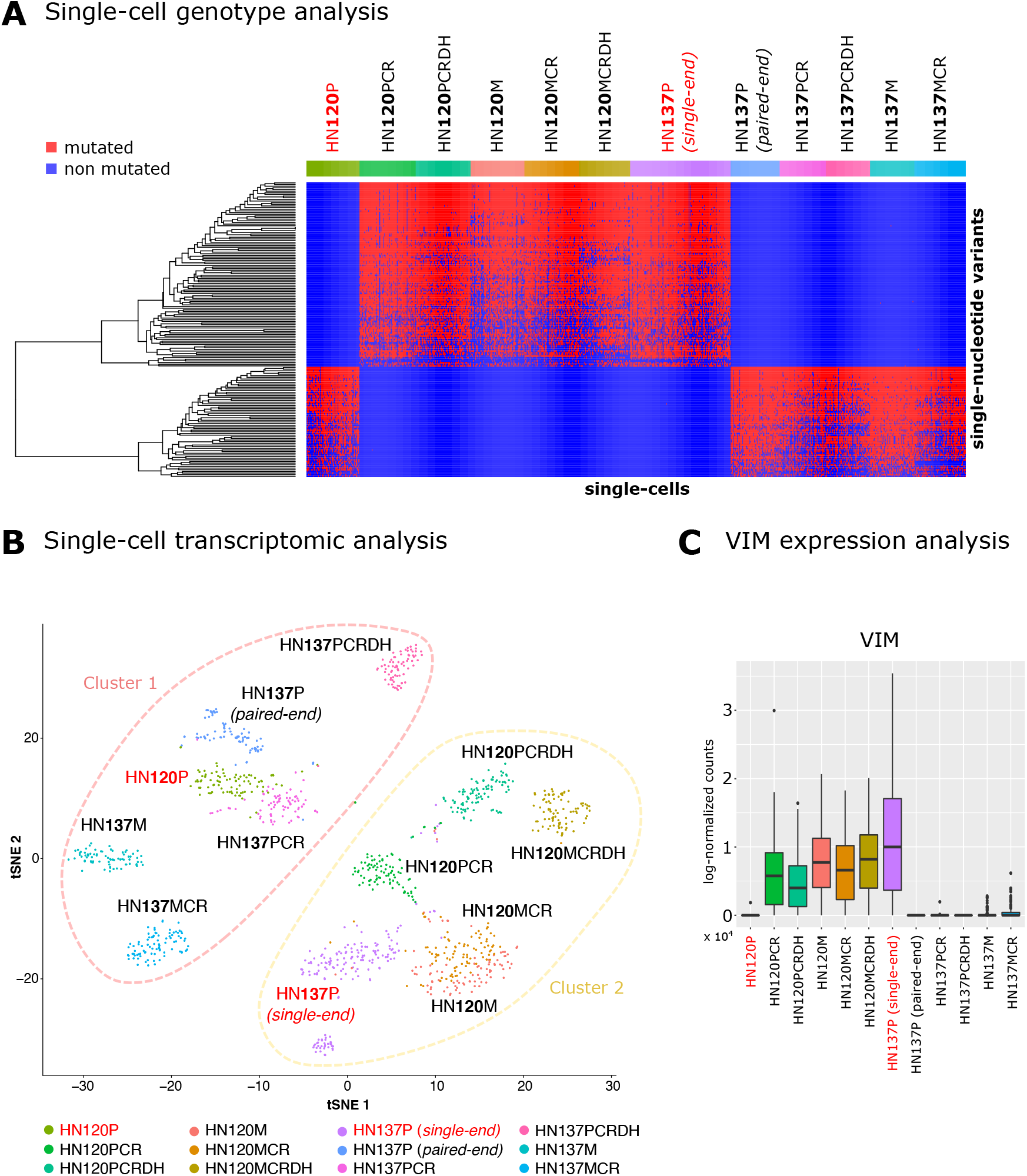
**(A)** The heatmap including the mutational profiles of all single cells of the HN120 and HN137 datasets is displayed (-P: primary line, -M: metastastic line, -CR: after cisplatin treatment, -CRDH: after drug-holiday). Red entries mark cells displaying a variant. For the ID of single-cells and SNVs please refer to Supplementary Table 1 and 2. **(B)** The t-SNE plot generated from the gene expression profiles of all single cells for all datasets is shown. **(C)** The distribution of the expression level of VIM on all single cells is shown with boxplots for all datasets.

Analogously, we identified 112 unique SNVs that are strongly informative for HN137P (*single-end*) identity, and that are present in 0 cells of HN120P (see Figure 1A). Such variants are observed in high frequency in HN137P (*single-end*) *and* in HN120PCR, HN120PCRDH, HN120M, HN120MCR, HN120MCRDH, whereas are not observed (< 1% of the cells) in HN137P (*paired-end*), HN137PCR, HN137PCRDH, HN137M, HN137MCR and HN120P. Supplementary Table 2 includes a summary of the analysis, in which for each SNV we report: genome position, reference and alternative alleles, rsID (if available), minor allele frequency (if available), the count and the ratio of single cells displaying the variant (total read count >= 5, alternative read count >= 3) in each dataset, the average total read count and the average alternative read count relative to the variant in each dataset.

From this analysis it is evident that the genotypic identity of HN120P cell line is inconsistent with that of the other HN120 datasets and with that of HN137P (*single-end*), whereas it is consistent with that of the remaining HN137 datasets. Conversely, the genotypic identity of HN137P (*single-end*) cell line is inconsistent with that of the other HN137 datasets and with that of HN120P, whereas it is consistent with that of all the other HN120 datasets. This consideration holds whether such SNVs are either germline or somatic, as genotypes are unquestionable footprints of cell identity (notice also that 177 on 179 variants have a rsID). These surprising results can be hardly explained by cancer-related selection phenomena, by random effects, or by sampling limitations. Instead, these observations suggest the presence of a methodological issue, which might be explained by a label swap of samples HN120P and HN137P (*single-end*).

This hypothesis is further supported by the single-cell transcriptomic analysis, which we performed via Seurat [15] (details are provided in the Supplementary Material). In Figure 1B one can find the t-SNE plot as computed on the 1000 most variable genes and in which single-cells are colored according to dataset label. Consistently with the genotype analysis, the transcriptomic analysis of single-cells highlights the presence of two distinct clusters, the first one including HN120P cells and all cells from HN137 datasets, excluded HN137P (*single-end*), the second one including HN137P (*single-end*) cells and all cells from HN120 datasets, excluded HN120P.

Unfortunately, we believe that this methodological error might have led to erroneous conclusions in [13, 14, 12]. In [13], for instance, the authors state that HN137 cell line is comprised of a mix of epithelial (ECAD+) and mesenchymal (VIM+) cells, whereas the HN120 cell line would include phenotypically homogeneous population of ECAD+cells. However, by looking at the expression level of VIM (Figure 1C), one can clearly notice that this gene is up-regulated in HN137P (*single-end*) and in all HN120 datasets, excluded HN120P, whereas is down-regulated (median = 0) in HN120P and in all HN137 datasets, excluded HN137(*single-end*).

Furthermore, in [13] the authors state that, in presence of cisplatin treatment, the heterogeneous HN137P cells demonstrate a progressive enrichment of ECAD, and the gradual depletion of VIM+cells, until the latter get extincted. Conversely, from the supposedly homogeneous ECAD+population of HN120P cells, the authors report the de novo emergence of VIM+cells after two weeks of treatment. In order to explain this unexpected phenomenon, the authors invoke the presence of a covert epigenetic mechanism that emerges under drug-induced selective pressure. Instead, we believe that this result might be easily explained by a label swap of HN120P and HN137P (*single-end*), as confirmed by the genotypic and transcriptomic analyses presented above (see Figure 1).

To conclude, the results presented in this work prove the efficacy of a joint analysis of the genotypic and transcriptomic identity of single-cells. Accordingly, this might represent a powerful instrument to uncover the elusive genotype-phenotype relation and to investigate the complex interplay underlying cancer evolution and drug resistance.

## Data availability

A repository including data and scripts to replicate the analyses is available at this link: https://github.com/BIMIB-DISCo/oral_squamous_longitudinal.

## Acknowledgements

This work was partially supported by the Elixir Italian Chapter and the SysBioNet project, a Ministero dell’Istruzione, dell’Università e della Ricerca initiative for the Italian Roadmap of European Strategy Forum on Research Infrastructures by the AIRC-IG grant 22082. Support was also provided by the CRUK/AIRC Accelerator Award #22790, “Single-cell Cancer Evolution in the Clinic”. We thank Giulio Caravagna, Chiara Damiani, Francesco Craighero and Lucrezia Patruno for helpful discussions.

## Contributions

All authors performed the analyses, interpreted the results, drafted and approved the manuscript. AG and DR supervised the study.

