## Supplementary Material for "Variant calling from scRNA-seq data allows the assessment of cellular identity in patient-derived cell lines"

**ARISING FROM** Sharma et al. *Nature Communication*
`https://doi.org/10.1038/s41467-018-07261-3`.

### GATK pipeline for variant calling from scRNA-seq data

To generate single-cell mutational profiles from scRNA-seq data, we employed the GATK Best Practices [2].

In particular, we selected the data uploaded on GEO repository with accession number GSE117872. As reported in the original work [6], single-cell RNA-seq were generated with the C1 Single-Cell Auto Prep IFC (Fluidigm) system. Single-cell libraries were prepared by using the Nextera DNA Sample Preparation Kit and the Nextera Index Kit (Illumina).

We downloaded the sample metadata by using the R library *GEOquery*. Single cells in the datasets were sequenced with two different library layouts and, in particular, all cell lines at each time point were sequenced using a *single-end* library layout, exception made for: $(i)$ the HN137M (metastatic cell line before treatment), which includes 75 single cells sequenced with paired-end library layout; $(ii)$ the HN137P (primary line before treatment), which includes 83 single cells sequenced with paired-end library layout and 170 with single-end library layout; moreover, 80 of such 170 cells were sequenced in two runs and are associated to two distinct FASTQ files.

We downloaded the RNA sequences using SRA toolkit. For each single cell sequenced in only one run with a single-end library layout, we obtained a single FASTQ file. For any single cell with double sequencing runs we concatenated the two FASTQ files. For cells with paired-end library layout, we only consider the forward reads files. As a result, we processed a single FASTQ file for each single cell of all datasets.

To summarize, the sample size for each dataset used in the analysis is the following: HN120P: 90 single cells, HN120PCR: 95, HN120PCRDH: 93, HN120M: 91, HN120MCR: 92, HN120MCRDH: 87, HN137P (*single-end*): 170 (80 of which with duplicated runs), HN137P (*paired-end*): 83, HN137PCR: 78, HN137PCRDH: 75, HN137M: 75, HN137MCR: 87.

By using Trimmomatic (v. 0.39) we removed the nucleotides with low quality score from the RNA sequences [1]. We then called SNVs and Indels in single cells by employing the GATK Best Practices. In particular, we aligned the single cell reads on the human reference genome (GRCh38 release) using the STAR aligner in 2-pass mode [3]. Then, we used Picard tools to preprocess the SAM files by adding read groups, sorting, marking duplicates and indexing. We used GATK (v. 3.8.1) to hard clip intronic regions with SplitNCigarReads utility and to re-calibrate base alignment by using BaseRecalibrator utility. This step requires the information about known single nucleotide polymorphisms (SNPs), which we retrieved on the dbSNP 1000 genome project phase 3. Finally, we used HaplotypeCaller and VariantFiltration to call genotype variants and filter out those with low quality score (with default parameters).

After applying the GATK pipeline, we generated a VCF file for each single cell. We used Annovar [8] to annotate the variants (i.e., synonymous or non-synonymous SNV, frame-shift insertion or deletion, stop-gain or stop-loss) and to add the rsID of each detected mutation, if available. Finally, we merged all the single-cell VCF files and applied custom filters as explained in the main text.

### Single-cell transcriptomic analysis via Seurat

Single-cell transcriptomic analysis was performed via Seurat [7]. In particular, we processed the normalized data included in the original dataset (GEO online repository, accession code `GSE117872`). Data were log-scaled and Z-score normalized. The 1000 most variable genes were then selected by using the variance stabilizing transformation (VST) [4]. We finally computed Principal Component Analysis on the selected genes and we employed the first 20 components to run the t-SNE algorithm for dimensionality reduction [5].
